# The intrinsic chaperone network of Arabidopsis stem cells confers protection against proteotoxic stress

**DOI:** 10.1101/2021.01.19.427268

**Authors:** Ernesto Llamas, Salvador Torres-Montilla, Hyun Ju Lee, María Victoria Barja, Elena Schlimgen, Nick Dunken, Prerana Wagle, Wolfgang Werr, Alga Zuccaro, Manuel Rodríguez-Concepción, David Vilchez

## Abstract

The biological purpose of plant stem cells is to maintain themselves while providing new pools of differentiated cells that form organs and rejuvenate or replace damaged tissues^1-3^. Protein homeostasis, or proteostasis, is required for cell function and viability^4-7^. However, the link between proteostasis and plant stem cell identity remains unknown. In contrast to their differentiated counterparts, we find that root stem cells can prevent the accumulation of aggregated proteins even under proteotoxic stress conditions such as heat stress or proteasome inhibition. Notably, root stem cells exhibit enhanced expression of distinct chaperones that maintain proteome integrity. Particularly, intrinsic high levels of the TRiC/CCT chaperonin determine stem cell maintenance and their remarkable ability to suppress protein aggregation. Overexpression of CCT8, a key activator of TRiC/CCT assembly^8^, is sufficient to ameliorate protein aggregation in differentiated cells and confer resistance to proteotoxic stress in plants. Taken together, our results indicate that enhanced proteostasis mechanisms in stem cells could be an important requirement for plants to persist under extreme environmental conditions and reach extreme long ages. Thus, proteostasis of stem cells could provide insights to design and breed plants tolerant to environmental challenges caused by the climate change.

## Introduction

Since proteins are involved in almost every biological process, protein homeostasis (proteostasis) is an essential requirement for cell physiology and viability. A complex network of cellular pathways maintains the proper concentration, folding, and interactions of proteins from their synthesis through their degradation. As such, the proteostasis network assures the integrity and quality of the proteome, preventing cell malfunction and death. However, aging as well as metabolic, environmental and pathological conditions can challenge the quality of the proteome in differentiated cells across tissues^4-7^.

Unlike most animals, plants exhibit a continuous supply of new and rejuvenated differentiated cells from the stem cell pools located in the root and shoot meristems^3^. For instance, the Sequoia tree contains stem cell reservoirs that can be active for more than 2,000 years^2^. As sessile organisms unable to remove themselves from persistent stressful environments^9^, this feature is particularly relevant to overcome everchanging conditions. Plant stem cells give rise to new organs or rejuvenate and repair tissues, allowing the plant to persist amidst variable and extreme conditions^1^. Thus, defining molecular and cellular differences between stem cells and their differentiated counterparts could shed light on how plants can live many years even under variable environmental stress conditions^1^. In these lines, recent findings demonstrate that the shoot meristems of 200 years-old oak trees are protected from the accumulation of deleterious mutations^1,10^. Given the essential role of proteostasis for cell function and viability, here we asked whether plant stem cells have enhanced proteostasis mechanisms to maintain their biological function.

## Results

To assess the proteostasis capacity of root stem cells of *Arabidopsis thaliana*, we used ProteoStat, a dye that becomes highly fluorescent when it binds to misfolded or aggregated proteins^11-13^. First, we performed a ProteoStat staining comparing seedlings grown under either control or distinct proteotoxic stress conditions, that is proteasome inhibition (*i.e.*, MG-132 treatment) and heat stress. In control seedlings, we did not detect aggregated proteins in most of the cells (**Fig. 1a and Supplementary Fig. 1**). The only exception was the cell population forming the sloughing lateral root cap, a layer that is continuously replaced through programmed cell death^14^ (**Fig. 1a and Supplementary Fig. 1**). On the other hand, the seedlings subjected to proteotoxic stress exhibited high levels of aggregated proteins across differentiated cells of the root (**Fig. 1a**) and cotyledons (**Supplementary Fig. 2a-b**).

**Fig. 1.**
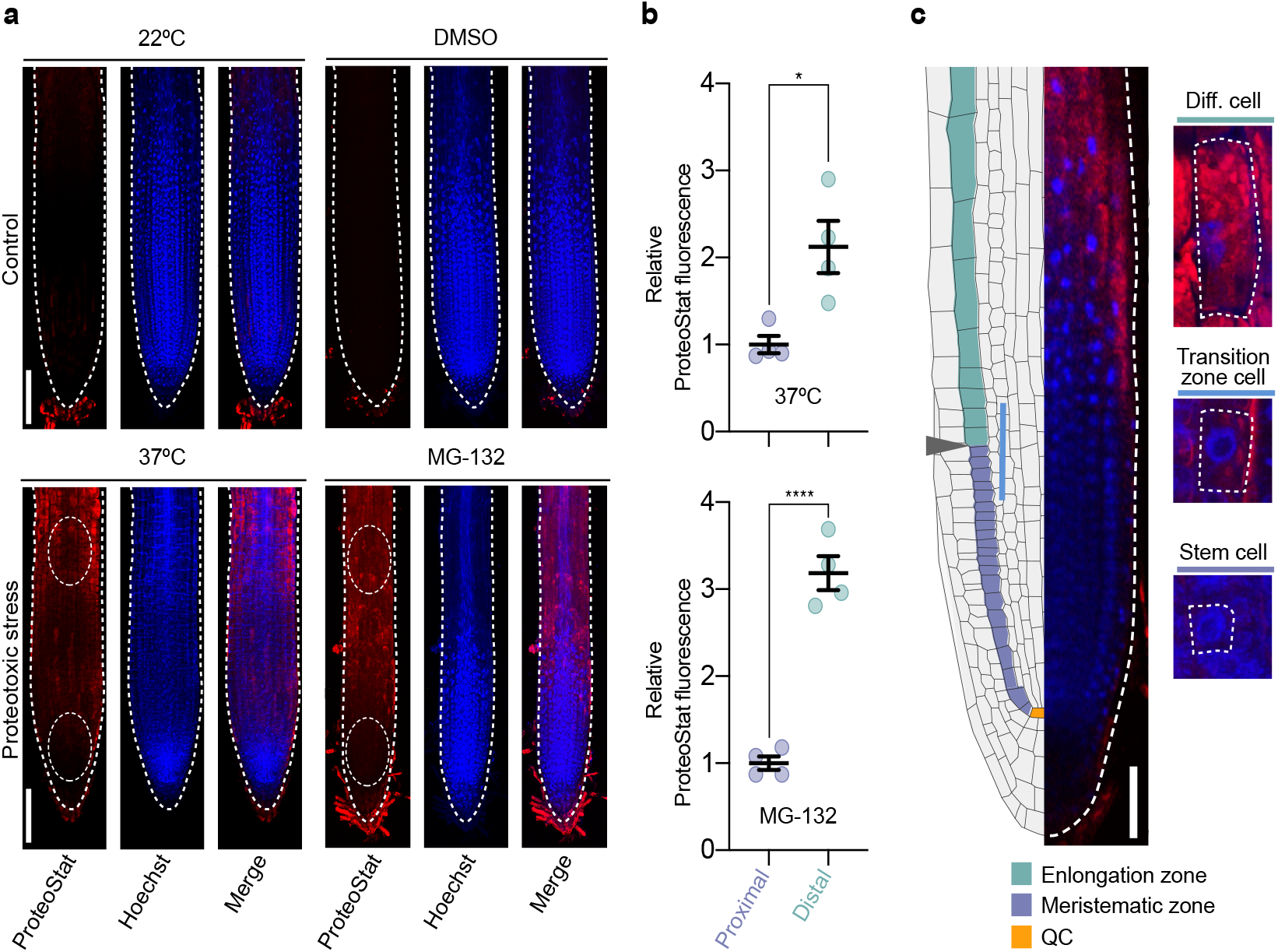
Proteotoxic stress causes differential protein aggregation in the distinct cell types of the root. **a,** Representative images of wild-type plant roots grown under control (22°C or 22°C + DMSO) or proteotoxic stress conditions stained with ProteoStat Aggregosome Detection kit. ProteoStat (red, protein aggregates), Hoechst (blue, nuclei), and Merge (ProteoStat and Hoechst) images are shown. For heat-stress assay, 4 days after germination (DAG) plants were transferred to 37°C for 2 days. For proteasome inhibition experiments, plants were germinated and grown in plates supplemented with 30 μM MG-132 and analyzed at 6 DAG stage. White bars represents 100 μm. **b,** Relative ProteoStat fluorescence levels comparing the area containing proximal (close to the QC) and distal cells from roots under proteotoxic stress (areas used form quantification are indicated with white dotted ovals in **Fig. 1a**). Scatter plots represent mean ± s.e.m of four independent experiments. The statistical comparisons were made by two-tailed Student’s *t*-test for unpaired samples. *P* values: \**P*<0.05, \*\*\*\**P*<0.0001. **c,** Left side, scheme indicating the structure of the Arabidopsis root meristem. The QC is indicated in orange color, the meristematic zone in green, and the elongation zone in purple. Gray arrowhead indicates the transition zone, where cells leave the meristem and enter the elongation/differentiation zone. Right side, close examination of a representative MG-132-treated root showing differential protein aggregation in cortical cells dividing and differentiating in opposite direction to the QC. White bar indicates 50 μm.

Damaged proteins in roots under stress showed a particular distribution, as the stem cells next to the quiescent centre (QC) had reduced amounts of protein aggregates compared with the rest of cells differentiating and expanding in opposite direction to the QC (**Fig. 1a-b**). A closer magnification indicated that misfolded and aggregated proteins can be gradually detected as the differentiation of cortical cells advances in opposite direction to the QC (**Fig. 1c**). Strikingly, cortical stem cells close to the QC showed undetectable protein aggregates (**Fig. 1c**). Thus, our data indicate that plant stem cells have an enhanced ability to prevent protein misfolding and aggregation under stress conditions compared with their differentiated counterparts. Stress conditions not only induced protein aggregation in differentiated cells, but also impaired growth (**Supplementary Fig. 3a-b**). However, plants recovered growth after removal of the stress, suggesting that stem cells were able to restart their function to provide new pools of differentiated cells after maintaining their proteostasis under stress (**Supplementary Fig. 4**).

The chaperome network is a key node of proteostasis to prevent protein misfolding and aggregation^15^. In Arabidopsis, the chaperome network is formed by more than 300 chaperones and co-chaperones that regulate protein folding and aggregation under normal and stress conditions^16^. To determine the molecular mechanisms that underlie the enhanced ability of stem cells to face proteotoxic stress, we performed a transcriptomic analysis of available RNA-seq data from two different pools of root cells isolated by fluorescence-activated cell sorting (FACS)^17^. By comparing root cells from the proximal part (mostly stem cells) and distal part (differentiated cells) respect to the QC (**Fig. 2a**), we identified differentially expressed components of the chaperome network such as distinct chaperones, co-chaperones and foldases in stem cells (**Fig. 2b**). Undifferentiated stem cells displayed more and larger-magnitude transcriptional changes than their differentiated distal counterparts (**Fig. 2b and Supplementary Data 1**). Indeed, almost half (49.4%) of the chaperome network was significantly up-regulated (*log*_2_ fold change >2; *P*<0.05) in stem cells (**Fig. 2b and Supplementary Data 1**). On the other hand, 35.9% of the chaperome was significantly downregulated (*log*_2_ fold change<-2; *P*<0.05) in these cells (**Fig. 2b and Supplementary Data 1**).

**Fig. 2.**
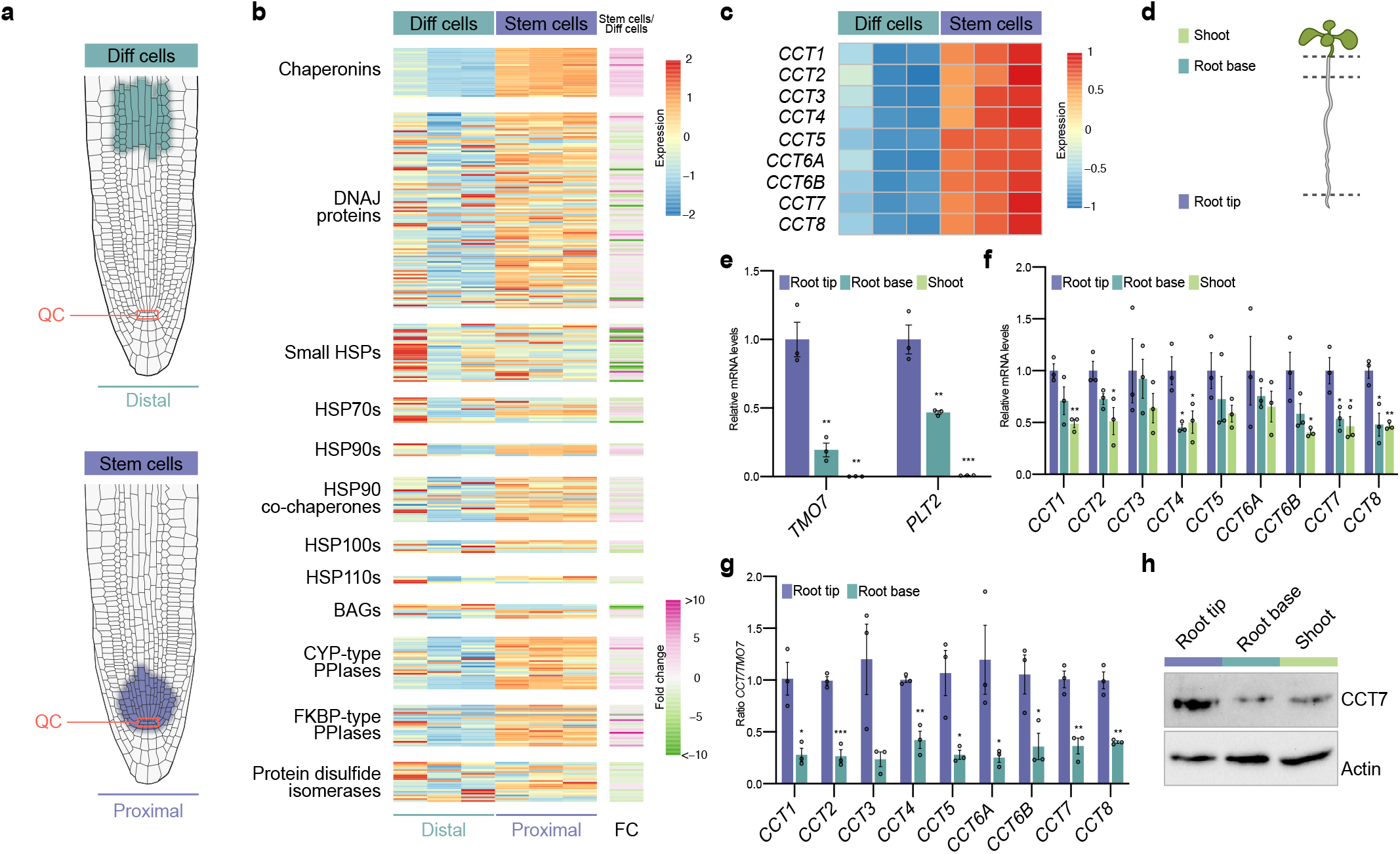
CCT subunits are upregulated in the root tip containing stem cells. **a,** Diagram depicting cell populations in the proximal zone (stem cells) and distal zone (differentiated cells) respect to the QC of the root apical meristem. Data for RNA-seq analysis were obtained from the indicated proximal and distal cells isolated by FACs (Wendrich *et al.*, 2017). **b,** Heatmap representing expression (fragments per kilobase of transcript per million, FKPM) and fold change expression of chaperome component transcripts comparing stem cells (proximal zone) with differentiated cells (distal zone) (*log*_2_ fold change (FC)>2, P<0.05 was considered significant). The chaperome genes were divided into different groups: Chaperonins (HSP60s), DNAJ proteins, small HSPs, HSP70s, HSP90s, HSP90 co-chaperones, HSP100s, HSP110s, BAGs, CYP-type PPIases, FKBP-type PPIases, and protein disulfide isomerase as defined by Finka *et al.*, 2011. **c,** Heatmap showing expression levels (FPKM) of all CCT subunits. Both FPKM heatmaps are row-scaled. **d,** Scheme depicting the different sections of wild-type plants at 6 DAG. Isolated shoot (green), root base (blue) and root tip (purple) were used for RNA and protein extraction. **e,** qPCR analysis of transcript levels of genes that are highly expressed in stem cells to regulate root meristem maintenance (mean ± s.e.m of 3 independent experiment). **f,** qPCR analysis of transcript levels of the indicated *CCT* subunits relative to levels in the root tip (mean ± s.e.m of 3 independent experiment). **g,** *CCT/TMO7* ratio in root tip and root bases samples (mean ± s.e.m of 3 independent experiment). **h,** Western blot analysis with antibody against CCT7 subunit. Actin is the loading control. The images are representative of three independent experiments. All the statistical comparisons between samples from root tip and differentiated cells were made by two-tailed Student’s *t*-test for unpaired samples. *P* value: \**P*<0.05, \*\**P*<0.01, \*\*\**P*<0.001.

Remarkably, our transcriptomic analysis revealed that most members (88.9%) of the heat shock protein (HSP) 60 family, also known as chaperonins, were significantly increased in root stem cells (**Fig. 2b and Supplementary Data 1**). Among them, all the subunits of the TRiC/CCT complex were upregulated in root stem cells (**Fig. 2c and Supplementary Data 1**). Since the TRiC/CCT complex is also highly expressed in mammalian embryonic and adult stem cells and it is critical for their function^8,18^, we focused on this specific node of the chaperome network. First, we assessed the increased expression of *CCT* subunits by quantitative RT-PCR (qPCR) analysis of samples manually isolated from the root tip containing mainly stem cells in comparison with the root base and the shoot, both of them containing mainly differentiated cells (**Fig. 2d**). The qPCR analysis of root meristem maintenance marker genes (*i.e., PTL2* (AT1G51190) and *TMO7* (AT1G74500)) confirmed that the distinct collected tissues contained different proportions of stem and differentiated cells (**Fig. 2e**). Both marker genes showed a gradient of expression with a maximum in the root stem cells^17,19^, as we observed in root tip samples (**Fig. 2e**). Likewise, qPCR analysis indicated that *CCT* transcripts are expressed at higher levels in the root tip compared with differentiated tissues (**Fig. 2f**). To further support increased expression of CCT subunits in stem cells, we calculated the ratio between individual *CCT* and *TMO7* transcripts levels obtained from each isolated root tissues, and we found that *CCT/TMO7* ratios were consistently higher in root tips compared with the root base (**Fig. 2g**). Importantly, western blot analysis confirmed that the higher mRNA levels of *CCT7* correlated with higher amounts of the protein in the root tip (**Fig. 2h**). In addition, the Plant eFP Viewer (bar.utoronto.ca), a bioinformatic tool that displays gene expression patterns, indicates higher expression levels of all the *CCT* genes in the root stem cell niche (**Supplementary Fig. 5a**). This analysis also shows enhanced expression of *CCT* genes in the meristematic zone enriched for stem cells of the xylem when compared with vascular cells of the distal xylem that have undergone a gradual differentiation process along the root (**Supplementary Fig. 5b**). Altogether, our data indicate that root plant stem cells have an intrinsic chaperome network characterized by enhanced expression of CCT subunits, a feature that could contribute to their enhanced ability to prevent protein aggregation under proteotoxic stress. Importantly, *CCT* transcripts were not upregulated in the root after heat stress treatment (**Supplementary Fig. 6**), suggesting that the basal high expression of CCTs is sufficient to prevent proteostasis collapse under stress conditions.

Intrigued by these results, we asked whether TRiC/CCT determines stem cell function and viability. Since loss of a single CCT subunit is sufficient to impair the assembly and activity of the TRiC/CCT complex^20-22^, we assessed whether an unbalance in the expression of CCT subunits impairs stem cell activity and subsequent root growth. To this end, we performed a root growth screening among Arabidopsis T-DNA mutants from genes encoding CCT subunits. We first isolated and characterized two insertion alleles for CCT4, CCT7 and CCT8 subunits (**Supplementary Fig. 7a**). T-DNA insertions in the promoter of *CCT4* (*cct4-1* and *cct4-2*) or *CCT7* (*cct7-1*) led to slightly reduced expression, whereas insertions in transcribed but untranslated regions of *CCT7* (*cct7-2*) and *CCT8* (*cct8-2* and *cct8-4*) caused a stronger reduction in transcript levels (**Supplementary Fig. 7b-c**). Thus, we classified the mutants as weak (*cct4-1, cct4-2* and *cct7-1*) and moderate loss of function (*cct7-2, cct8-2* and *cct8-4*). While we did not observe root growth effects in weak mutants, all the moderate mutant lines exhibited shorter roots compared with wild-type (WT) controls (**Supplementary Fig. 8**). In addition, we also examined the root length of mutants with alterations in the prefoldin (PFD) complex^23^, a distinct molecular chaperonin which is also highly expressed in stem cells (**Fig. 2b and Supplementary Data 1**). Importantly, PFD and TRiC/CCT complex interact and act together to facilitate folding of numerous proteins^24^. Similar to moderate *cct7-2, cct8-2* and *cct8-4* mutants, *pfd3* and *pfd5* mutants showed shorter roots compared with the WT seedlings (**Supplementary Fig. 8**). Thus, our data indicate that TRiC/CCT along with PFD complex contribute to proper maintenance of the meristem and subsequent continuous root growth.

Given that null mutations of CCT subunits are lethal in plants, yeast and mammals^25,26^, the moderate loss-of function mutants *cct7-2* and *cct8-2* provide an invaluable mean to study proteostasis and stem cell maintenance. We further characterized these *cct* mutants and found that the root growth was impaired over time (**Fig. 3a**). Moreover, mutations in *CCT* genes lead to a reduced number of cortical cells in the meristem, resulting in shorter meristems (**Fig. 3b-c**). Notably, *cct7-2* and *cct8-2* mutant plants displayed a disordered stem cell niche with aberrant cellular organization and division planes in the columella root cap and the QC (**Fig. 3c**). Prompted by these results, we tested whether downregulation of *CCT* levels alter expression of markers of stem cell maintenance. To this end, we compared the transcript levels of *PLT2* and *TMO7* between WT, *cct7-2* and *cct8-2* plants. Notably, we found lower levels of *PLT2* and *TMO7* in the *cct* mutant seedlings (**Supplementary Fig. 9**).

**Fig. 3.**
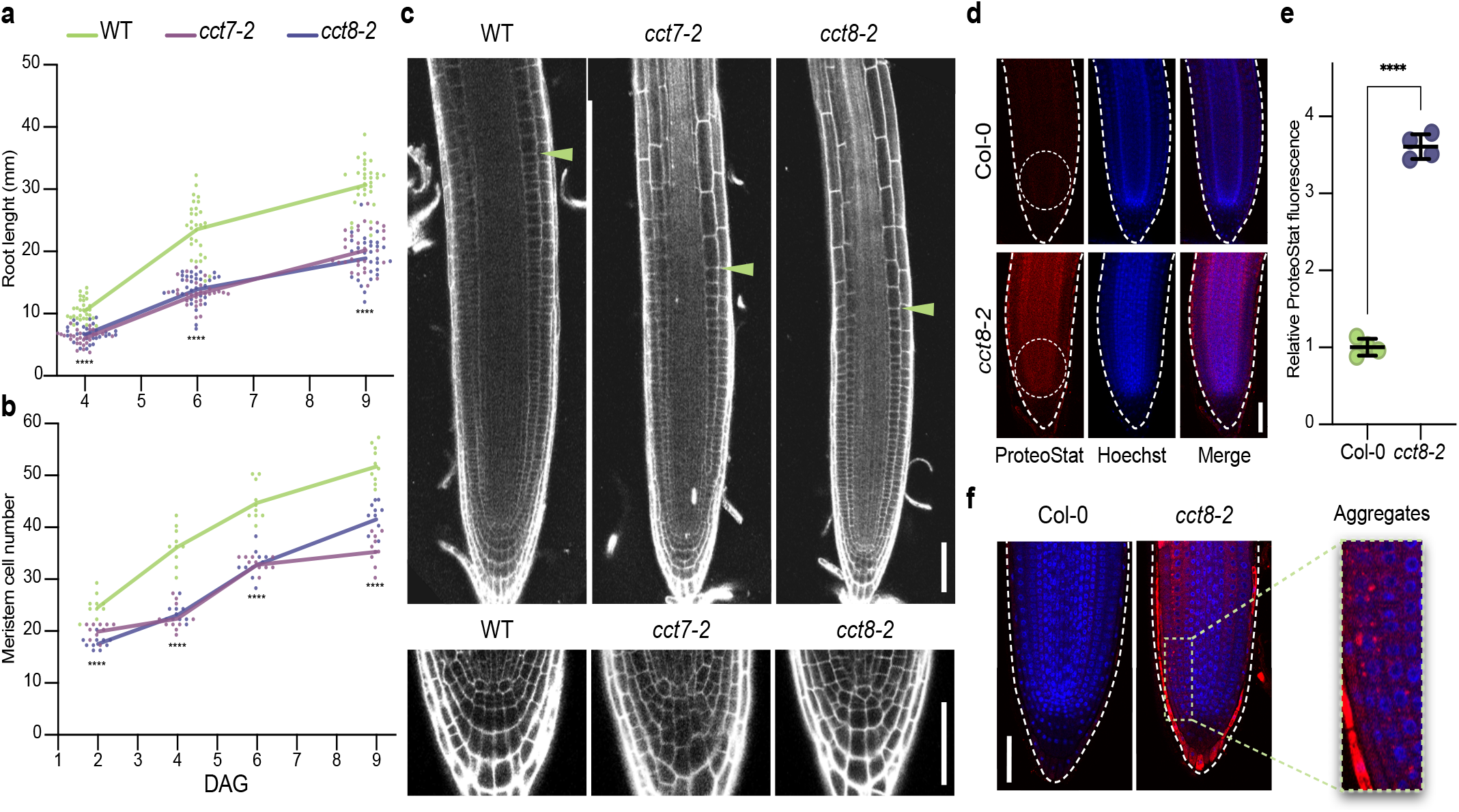
Reduced expression of CCT triggers protein aggregation in root stem cells and impairs stem cell function. **a,** Root length over time in wild-type (WT), *cct7-2*, and *cct8-2*. DAG, days after germination (n= 30 roots from three independent experiments). **b,** Meristem cell number over time in WT, *cct7-2*, and *cct8-2* (n= 10 roots from three independent experiments). In **a-b**, staggered plot showing connecting lines to the means. **c,** Representative images of propidium iodide (PI)-stained roots show differences between the meristem size of Col-0 WT and *cct* roots at 4 DAG. Green arrowheads indicate the junction between the meristematic and elongation zone. Lower panels show a magnification to the QC and columella cells. Scale bars represent 50 μm. **d,** Representative images of ProteoStat staining of Col-0 WT and *cct8-2* mutant at 6 DAG. Seedlings were grown under control conditions (22°C). **e,** Relative ProteoStat fluorescence levels within the area containing proximal cells (close to the QC) of *cct8-2* relative to Col-0 control (mean ± s.e.m of 4 independent experiments). The areas used for quantification are indicated with white dotted ovals in Fig. 3d. **f,** Higher magnification of the stem cell niches of Col-0 and *cct8-2*. All the statistical comparisons were made by two-tailed Student’s *t*-test for unpaired samples. *P* value: \*\*\*\**P*<0.0001.

We hypothesized that the imbalance in the TRiC/CCT complex of *cct* mutants could lead to aberrant protein aggregation in stem cells, a process that impairs cell function. To assess this possibility, we used ProteoStat staining. Notably, we found increased protein aggregation in the stem cell niche of *cct* mutant plants compared with WT controls (**Fig. 3d-e**). Moreover, a closer examination of the cortical stem cell region allowed us to detect fluorescence speckles in stem cells, indicating protein aggregation (**Fig. 3f**). Taken together, these data establish a link between proteostasis, TRiC/CCT complex activity and root stem cell maintenance.

During organismal aging, differentiated cells of animals undergo a progressive decline in their proteostasis network, losing their ability to maintain proteome integrity and cope with proteotoxic stress^27,28^. However, mammalian embryonic stem cells rely on enhanced proteostasis mechanisms to replicate indefinitely while maintaining their undifferentiated state and, therefore, are immortal in culture^8,29,30^. Since our results indicate that plant stem cells also exhibit enhanced proteostasis, we asked whether mimicking root stem cell proteostasis in somatic tissues confers organismal protection to proteotoxic stress. For this purpose, we overexpressed CCT8, a subunit that it is sufficient to increase TRiC/CCT assembly in both human cells and the roundworm *Caenorhabditis elegans*^8^. We generated two independent transgenic lines that express upregulated levels of CCT8 (**Supplementary Fig. 10**) and assessed resistance to proteotoxic stress. Notably, we found that CCT8-overexpressing plants accumulate less aggregates in roots compared with wild-type plants when both were treated with MG-132 proteasome inhibitor (**Fig. 4a-b**), resulting in longer roots under this deleterious condition (**Fig. 4c**). In addition, stress-challenged transgenic lines displayed extra cell layers in the sloughing lateral root cap that is normally subject to continuous cell death and replacement (**Supplementary Fig. 11a**). This phenotype could be caused by the constitutive CCT8 expression during the columella differentiation process, in which the levels of endogenous *CCT8* transcript decay (**Supplementary Fig. 11b-c**). Besides proteasome inhibition, we also assessed the effects on heat stress in the shoot of CCT8-overexpressing plants. To this end, we performed a heat-shock survival assay where 6 day after germination (DAG) seedlings were subjected to 3 hours of 45°C heat-shock and then shifted back to 22°C. We found that *35S:CCT8* plants exhibit increased survival after heat shock compared with wild-type plants (**Fig. 4d**). Since heat shock triggers the accumulation of misfolded and aggregated proteins that can cause cell death, enhanced survival after heat shock may be explained by reduced protein aggregation in *35S:CCT8* plants. In support of this hypothesis, soluble and insoluble protein analysis as well as ProteoStat staining confirmed that *35S:CCT8* seedlings contained less protein aggregates after heat shock (**Fig. 4e-g**). By native gel analysis, we confirmed that TRiC/CCT complex assembly was enhanced in plants upon CCT8 overexpression (**Fig. 4h**). Thus, our data indicate that upregulation of CCT8 and subsequent assembly of TRiC/ CCT complex confers protection to heat stress by sustaining the integrity of the proteome.

**Fig. 4.**
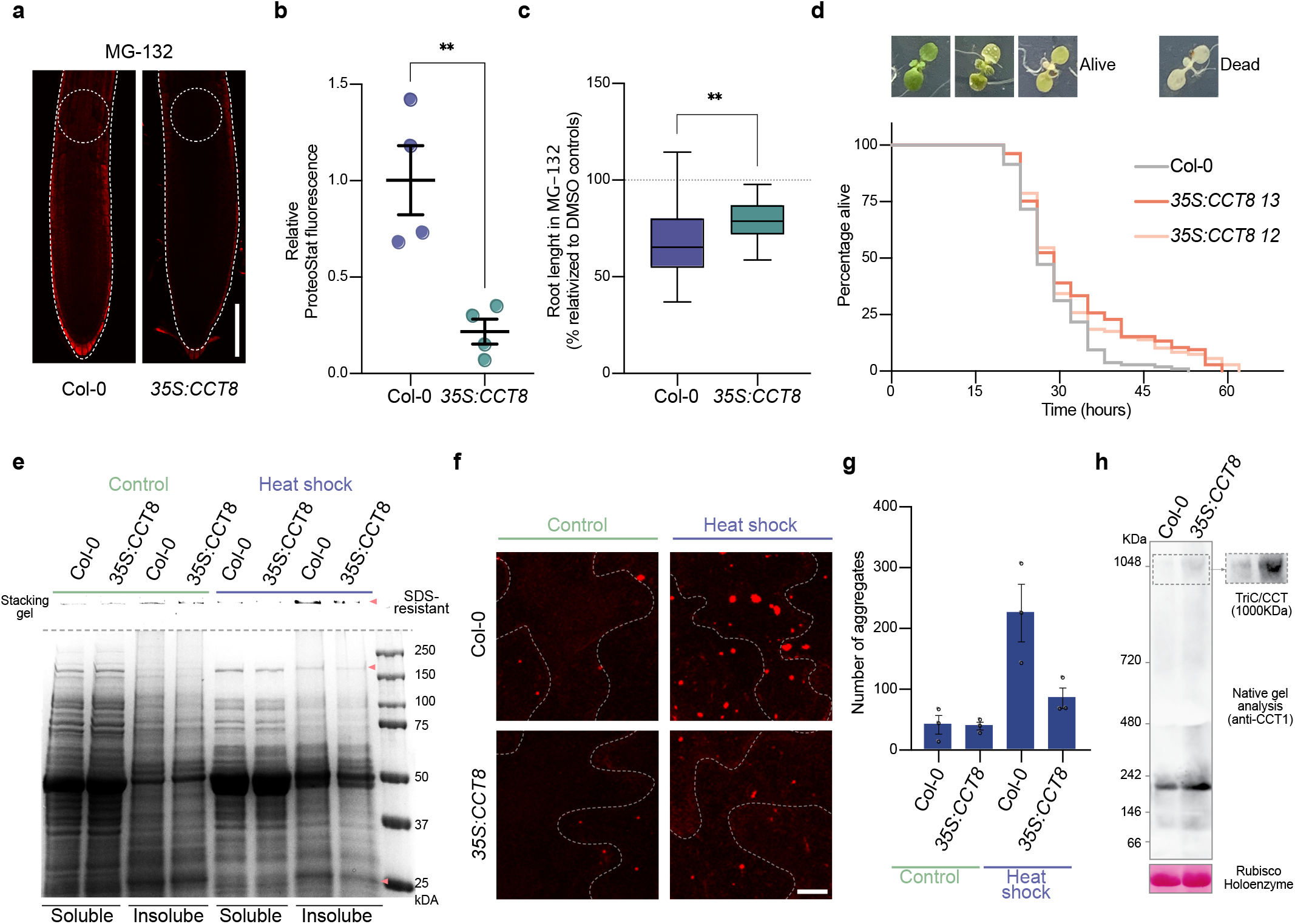
Overexpression of CCT8 is sufficient to ameliorate protein aggregation in differentiated cells and confer resistance to proteotoxic stress in plants. **a,** Representative images of ProteoStat staining of 6 DAG Col-0 and *35S:CCT8* germinated and grown in plates supplemented with 15 μM MG-132. Scale bar represents 100 μm. **b,** Relative ProteoStat fluorescence levels comparing the areas depicted with white dotted circles in **Fig. 4a**. Graph represents mean ± s.e.m of four independent experiments. **c,** Root lengths of MG-132-treated plants relative to their corresponding controls treated with DMSO. Statistical comparisons in b-c were made by two-tailed Student’s *t*-test for unpaired samples. *P* value: \*\**P*<0.01. **d,** Heat shock survival assay of *35S:CCT8* plants and Col-0 WT. 6 DAG seedlings grown at 22 °C were subject to 45°C for 3 h (heat shock) and then shifted back to control conditions (22°C). Alive plants were scored. Death plants were counted when they showed a complete pigment bleaching phenotype. Kaplan-Meier survival data is representative of 2 independent experiments. All lines were grown in the same plate. *35S:CCT8 12* (long rank, *p*<0.00138) and *35S:CCT8 13* (long rank, *p*<0.00151) survive longer than control seedlings under heat stress. Col-0 WT: median=26, n=76/108, *35S:CCT8 12:* median=29, n=59/108. *35S:CCT8 13:* median=29. **e,** Representative BlueSafe-stained SDS-PAGE gel of protein extracts from the soluble and insoluble fractions. Insoluble/aggregated protein bands that show strong decreased in heat stress-treated *35S:CCT8* plants compared with heat stress-treated WT are indicated with orange arrowheads. SDS-resistant aggregates retained on the stacking gel are also shown. **f,** Plants from the heat shock assay were stained with ProteoStat and representative images are shown. Scale bar represents 10 μm. **g,** Quantification of the number of aggregates stained with ProteoStat. Total number of aggregates were counted in the whole 63X captured image from the epidermis of the cotyledons (mean ± s.e.m from 3 independent experiments). **h,** Representative images of two independent native gel electrophoresis of Col-0 WT and *35S:CCT8* extracts collected from 14 DAG plants followed by immunobloting with CCT1 antibody (right panel shows the TRiC/CCT complex band with longer exposure time). Band corresponding to Rubisco holoenzyme after Ponceau staining is the loading control.

## Discussion

The cells of all living organisms contain numerous proteins which are at risk of misfolding and aggregation^31,32^. The accumulation of damaged proteins can lead to defects in growth, decrease in yield, accelerated aging, and cellular death^33^. A series of stringent proteostasis mechanisms maintain protein homeostasis^4,5^. However, aging, metabolic conditions and environmental challenges can overwhelm such proteostasis mechanisms. For example, protein aggregation in aging neurons is linked with several human neurodegenerative disorders such as Alzheimer’s, Parkinson’s or Huntington’s disease^27,34,35^. Like-wise, stress conditions can induce proteostasis collapse and protein aggregation in plants^12,36,37^. We found that maintaining low levels of protein aggregation during stress conditions contribute to stem cell viability, ensuring cellular replacement as well as continuous and repetitive formation of new structures and organs. Notably, root stem cells have an enhanced protein folding capacity, providing them with a striking ability to cope with proteotoxic stress when compared with their differentiated counterparts. Thus, both stem cells of plants and animals rely on a powerful proteostasis machinery for their function and maintenance^8,11,18,29,30,34,38-40^.

Plant and animal stem cells have unexpected similarities that modulate their function^41^. One classic paradigm is the conserved protein Retinoblastoma (Rb), which is critical for cell division and differentiation of plant and animal stem cells^2,41^. In addition, genome stability maintains a healthy pool of stem cells in both plants and animals^10,42^. Recently, it has been reported that stem cells of the shoot meristem are robustly protected from the accumulation of mutations in long-lived trees^10^. Besides cell division and genome stability, our results establish that both plant and animal stem cells share an enhanced proteostasis network. For instance, similar to animal stem cells^8,18,43^, we identified distinct components of the Arabidopsis chaperome upregulated in stem cells, such as DNAJs and HSPs. Particularly, we found a strong enrichment in the levels of the distinct members of the chaperonin family in stem cells, including all the CCT subunits of the TRiC/CCT complex. Preventing protein misfolding and aggregation in plant stem cells before, during and after stress conditions can be an important requirement for stem cell viability and function. As such, the super-vigilant proteostasis network of stem cells could allow the plant to keep growing during stress or regenerate after removal of the stress. In these lines, we speculate that the stringent proteostasis of plant stem cells contribute to the extreme ages that distinct plants are able to achieve, even when they are continuously exposed to abiotic stresses such as high temperature^3^. While very few mammals, including humans, can live more than hundred years, plants like the bristlecone pine and the giant sequoia are able to live over a thousand years^1,2,9^ probably due to their stem cell pools^9^.

Remarkably, we found that mimicking the proteostasis network of stem cells in somatic tissues by increasing TRiC/CCT assembly is sufficient to prevent protein aggregation in differentiated cells and increased organismal survival under stress conditions. Thus, rewiring proteostasis mechanisms in crop plants could have an important agronomic value to maintain yield in the face of unpredictable daily temperature fluctuations due to the global climate change.

## Methods

### Plant material, constructs and growth conditions

All the Arabidopsis thaliana lines used in this work are in Columbia-0 (Col-0) ecotype. Wild-type, and loss-of-function mutants *cct4-1* (SALKSEQ_069998.1), *cct4-2* (SALKSEQ_076214.0), *cct7-1* (SALKSEQ_135744.2), *cct7-2* (SALK_099986), *cct8-2* (SALK-SEQ_082168)^25^, *cct8-4* (SALKSEQ_137802), *pfd3*^44^ and *pfd5*^44^ were used in this study. T-DNA mutants were genotyped by PCR. All seeds were surface-sterilized and germinated on solid 0.5X Murashige and Skoog (MS) medium without sucrose neither vitamins, and plants were incubated in a growth chamber at 22°C (or otherwise indicated in the figure) under long-day conditions as previously described^45^. When indicated in the figure, medium was supplemented with MG-132 (Sigma). For the root growth analysis, we used the software MyRoot to semi-automatically measure the root length from 6 day after germination (DAG) seedlings grew in vertical agar plates^46^. For qPCR and western blot analysis of the root tip, base root and shoot, seeds were germinated on vertical plates with a sterile mesh (SefarNitex 03-100/44) placed on the medium. 6 DAG plants were dissected manually with a surgical blade for posterior RNA and protein extraction. For the generation of the *35S:CCT8* lines, the full-length coding sequence of Arabidopsis *CCT8* (AT3G03960) was cloned under the *35S* promoter using the entry vector pDONR207 and the destination vector pGWB502. Arabidopsis plants were transformed by the floral dip method^47^.

### Analysis of RNA-seq data

We analyzed public available RNA-Seq data from two different pool of cells isolated by FACS from Arabidopsis roots of the lines *PUB25* and *SPT*. We used the RNA-sequencing samples *PUB25* proximal 1-3 (corresponding to mainly stem cells) and *SPT* distal 1-3 (corresponding to differentiated vascular cells)^17^ to compare the expression profile of the Arabidopsis chaperome as defined by Finka *et al.* 2011. RNA-seq data were analysed using a QuickNGS pipeline^48^. This workflow system provided a basic read quality check using FastQC (version 0.10.1) and read statistics using SAMtools (version 0.1.19). The basic data processing of the Quick-NGS pipeline consists of a splicing-aware alignment using Tophat2 (version 2.0.10) followed by reference-guided transcriptome reassembly with Cufflinks2 (version 2.1.1). The QuickNGS pipeline calculated read count means, fold change and P values with DEseq2 and gene expression for the individual samples with Cufflinks2 (version 2.1.1) as fragments per kilobase of transcript per million (FPKMs), in both cases using genomic annotation from the Ensembl database, version 32. All data preprocessing and visualization was done with R version 3.2.2 and Bioconductor version 3.0. Differential expressed genes (DEGs) calculated by DESeq2 were filtered using P value <0.05 cutoff of significance, and *log*_2_ fold change>2 and *log*_2_ fold change<-2 cutoffs of relative expression for upregulated and downregulated DEGs respectively. Heatmaps were created using “pheatmap” package.

### Western blot analysis

Total protein extracts were obtained from Arabidopsis lyophilized powder. The powder was resuspended on ice-cold TKMES homogenization buffer (100 mM Tricine-potassium hydroxide pH 7.5, 10 mM KCl, 1 mM MgCl_2_, 1 mM EDTA, and 10% [w/v] Sucrose) supplemented with 0.2% (v/v) Triton X-100, 1 mM DTT, 100 μg/ml PMSF, 3 μg/ml E64, and 1X plant protease inhibitor (Sigma). The resuspended sample was centrifuged at 10,000xg for 10 min at 4°C and the supernatant recovered for a second step of centrifugation. Protein concentration was determined using the kit Pierce Coomassie Plus (Bradford) Protein-Assay (Thermo Scientific). Approximately 40-50 μg of total protein was separated by SDS-PAGE, transferred to nitrocellulose membrane and subjected to immunoblotting. The following antibodies were used: anti-CCT7 [1:1,000] (Abcam, ab170861) and anti-Actin [1:5000] (Agrisera, AS132640).

### Separation of soluble and insoluble protein extracts

For the separation of soluble and insoluble (with protein aggregates) fractions, native protein extracts were obtained using a buffer containing 100 mM Tris-HCl pH 7.9, 10 mM MgCl_2_, 1% (v/v) glycerol, and 1X plant protease inhibitor (Sigma). After centrifugation for 10 min at 10,000xg, the supernatant was collected as the soluble fraction. The pellet was washed with fresh buffer and centrifuged again. The obtained pellet fraction was then resuspended in denaturing TKMES buffer and centrifuged again to collect the supernatant as the insoluble fraction. Protein concentration from each fraction was determined with Pierce Coomassie Plus (Bradford) Protein-Assay (Thermo Scientific).

### Blue native gel immunoblotting of TRiC/CCT complex

Whole Arabidopsis seedlings were frozen in liquid nitrogen and ground to powder using a mortar and pestle. The frozen powder was collected in lysis buffer (50 mM Tris-HCl (pH 7.5), 1 mM dithiothreitol and 10% glycerol supplemented with 1X plant protease inhibitor (Sigma)). After centrifugation 10,000xg for 10 min at 4°C, the supernatant was collected and protein concentration was determined. 90 μg of total protein was run on a 3-13% gel in deep blue cathode buffer (50 mM Tricine, 7.5 mM Imidazole and 0.02% Coomassie G250) at 4°C for 3 h at 100 V and then exchange deep blue cathode buffer to slightly blue cathode buffer (50 mM Tricine, 7.5 mM Imidazole, and 0.002% Coomassie G250) and run at 100 V overnight. Proteins were then transferred to a polyvinylidene difluoride membrane at 400 mV for 3 h by semi-dry blotting. For loading control, the membrane was stained with Ponceau S. Western blot analysis was performed with a monoclonal antibody against CCT1 [1:1,000] (Abcam, ab109126).

### Gene expression analysis

Total RNA was extracted from plant tissue, using the Maxwell 16 LEV Plant RNA Kit (Promega). RNA was quantified using a NanoDrop (Thermo Scientific). cDNA was synthetized using the kit NZY First-Strand cDNA Synthesis Kit (nzytech). SybrGreen real-time quantitative PCR experiments were performed with a 1:20 dilution of cDNA using a CFC384 Real-Time System (Bio-Rad). Data were analyzed with the comparative 2ΔΔCt method using the geometric mean of *Ef1α* and *PP2A* as housekeeping genes. See **Supplementary Table 1** for qPCR primers used in this work.

### Confocal microscopy and fluorescence quantification

Confocal microscopy images were taken either with FV1000 Confocal Laser-scanning Microscope (Olympus) or a Meta 710 Confocal Microscope with laser ablation 266 nm (Zeiss). All images were acquired using the same parameters between experiments. For the detection of aggregated proteins, we used the ProteoStat Aggresome detection kit (Enzo Life Sciences). Seedlings were stained according to the manufacturer’s instructions. Briefly, seedlings were collected and were fixed in 4% formaldehyde solution for 30 min at room temperature. Formaldehyde solution was removed and seedlings were washed twice with 1X PBS. Then, seedlings were incubated with permeabilizing solution (0.5% Triton X-100, 3 mM EDTA, pH 8.0) with gently shaking for 30 min at 4°C. Seedlings were washed twice with 1X PBS. Then, plants were incubated with 1X PBS supplemented with 1 μL/mL of ProteoStat and 1 μL/mL Hoechst 33342 (nuclear staining) for 30 min at room temperature. Finally, seedlings were washed twice with 1X PBS and mounted on a slide. Quantification of ProteoStat fluorescence was performed with ImageJ software.

### Heat-shock survival assay

A single plate containing 6 DAG seedlings (control and CCT8 overexpressor lines) were covered with aluminum foil and transferred to 45°C for 3 h. Then, the plate was transferred back to 22°C under long-day conditions. 108 seedlings were used per line and scored every 3 h. Green seedlings (containing chlorophyll pigments) were considered alive while complete bleached seedlings were considered as death. PRISM 9 software was used for statistical analysis and P values were calculated using the *log*-rank (Mantel-Cox) method.

## Acknowledgments

This work was supported by the European Research Council (ERC Starting Grant-677427 StemProteostasis) and the Deutsche Forschungsgemeinschaft (DFG) (Germany’s Excellence Strategy-CECAD, EXC 2030-390661388). This work was also supported by the Humboldt Research Fellowship for postdoctoral researchers to E. Llamas. We thank the technical support from M.R. Rodríguez-Goberna, M. Chmara, and P. Comelli. We thank P. Mammadova and O. Waheed for helping with experiments. We also thank I. Betegón-Putze for helping with root analysis and measurement. We are grateful to J. Salinas for providing the *pfd3* and *pfd5* seeds. Anti-Actin antibody was a gift from J.F. Martínez-García. pGWB502 was a gift from T Nakagawa (Addgene plasmid 74844). We thank the Salk Institute Genomic Analysis Laboratory and NASC for providing the sequence indexed Arabidopsis T-DNA insertion mutants. We thank the CRAG (M. Amenós) and CECAD (A. Schauss, C. Jungst) imaging facilities for their support in confocal microscopy.

