## Supplementary Figures for "The intrinsic chaperone network of Arabidopsis stem cells confers protection against proteotoxic stress"

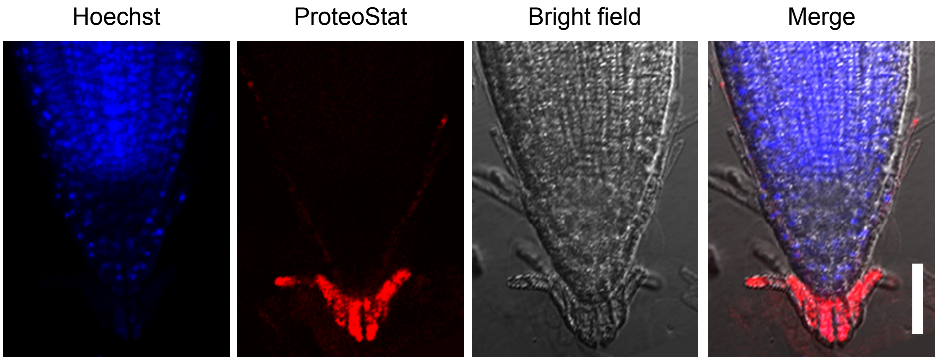


**Supplementary Figure 1. The cells of the sloughing lateral root cap exhibit high levels of protein aggregates under non-proteotoxic control conditions.** Representative images of a 6 DAG wild-type root grown at 22ºC on plates supplemented with DMSO and stained with ProteoStat. Scale bar indicates 50 µm.

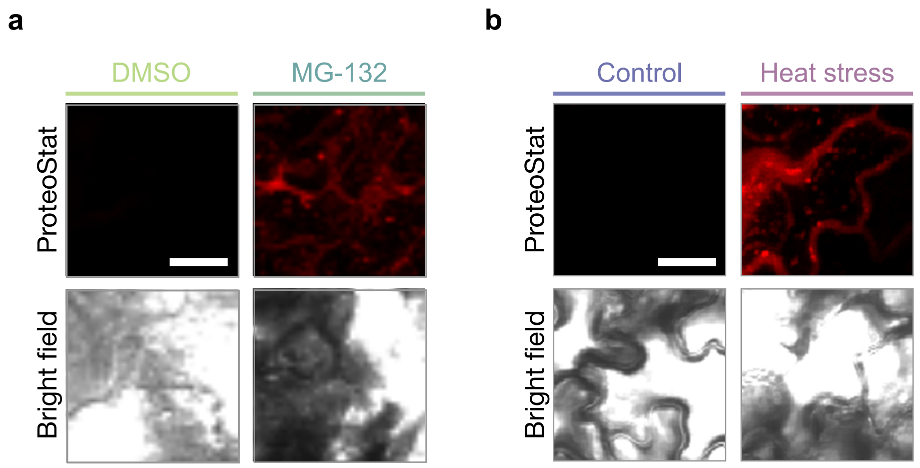


**Supplementary Figure 2. Proteotoxic stress induces protein aggregation in cotyledons.** Representative images of cotyledons of plants stained with ProteoStat upon 30 μM MG-132 treatment (**a**) or heat stress (**b**). Scale bars represent 20 μm.

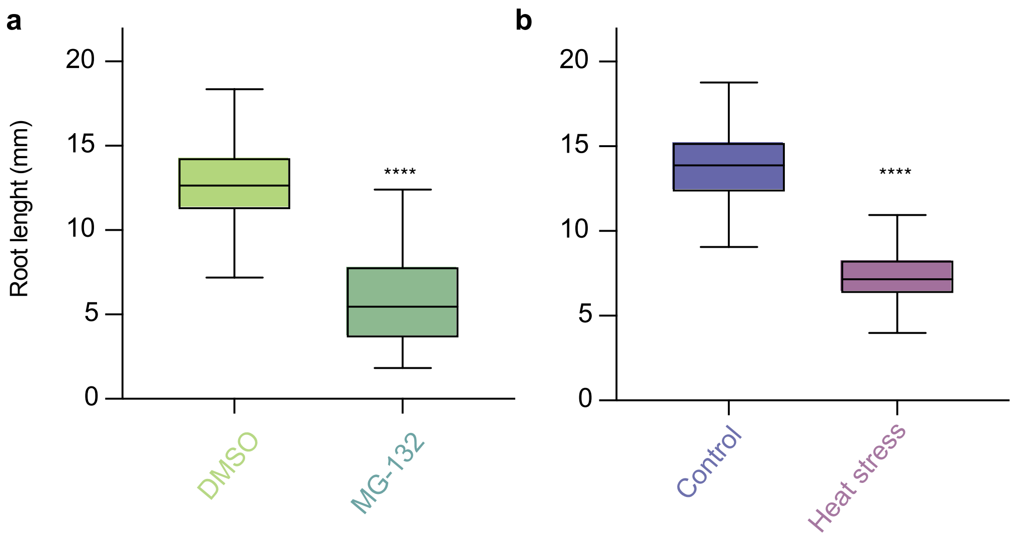


**Supplementary Figure 3. Proteotoxic stress decreases root growth. a,** Tukey box plots representing root length in millimeters (mm) of 6 DAG Col-0 wild-type plants grown on either DMSO or 30 µM MG-132 proteasome inhibitor. **b,** Root length of 6 DAG plants grown at 22ºC (Control) and 6 DAG plants that were transferred to 37 ºC at 4 DAG (Heat stress). Horizontal line in Tukey box plots represents the median and box represents 25th and 75th percentiles. Data from 3 independent experiments were analyzed. The statistical comparisons were made by two-tailed Student's *t*-test for unpaired samples. *P* value: ****$P<0.0001$.

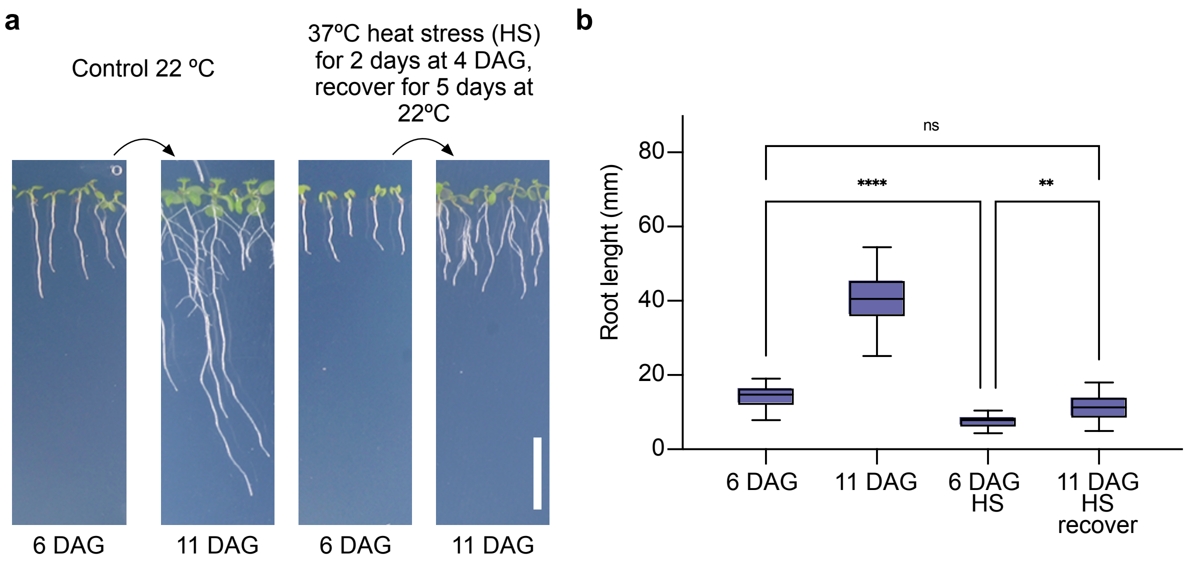


**Supplementary Figure 4. Plants recover growth after removal of proteotoxic stress. a,** Representative images of 6 DAG or 11 DAG wild-type seedlings grown continuously under control conditions (22ºC under long-day conditions) or plants treated with 2 days 37ºC heat stress (HS) when they were at 4 DAG stage (grown at 22ºC). At 6 DAG heat-treated plants were transfer back to recover at 22 ºC for 5 days under long-day conditions. DAG= days after germination. Scale bar represents 10 mm. **b,** Measurement of the root length under the conditions indicated above. Horizontal line in Tukey box plots represents the median and the box represents 25th and 75th percentiles (n= 30). Statistical analysis were performed with one-way ANOVA for multiple comparisons. *P* values: ***P*<0.01, *****P*<0.0001, ns= not significant (*P*>0.05).

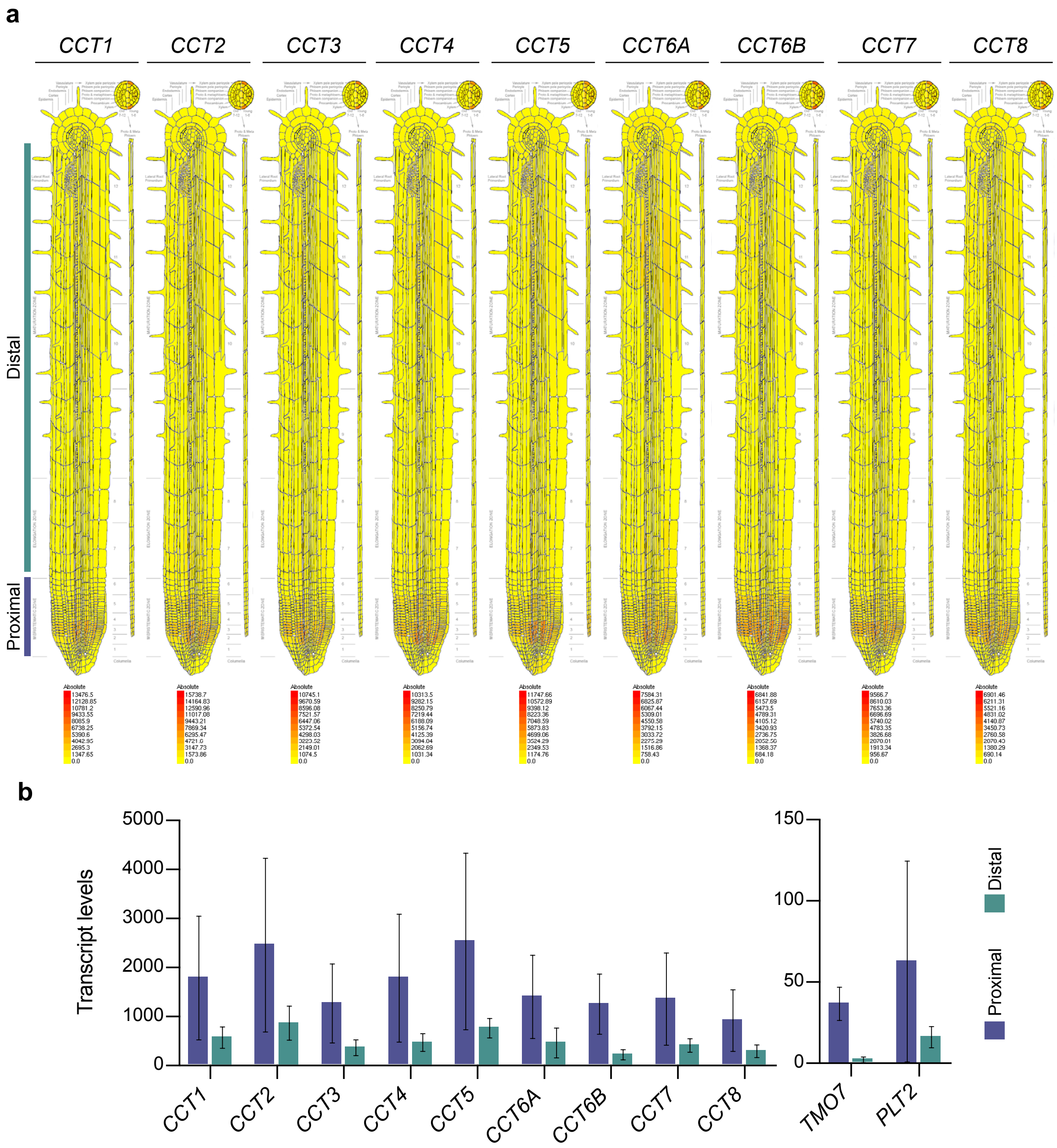


**Supplementary Figure 5. The root apical meristem exhibits high expression of *CCT* subunits. a,** Root expression maps indicating the spatial level of expression of the distinct Arabidopsis CCT subunits. **b,** Bar charts represent the expression levels of CCT subunits in xylem corresponding to proximal/young cells (meristematic zone) or distal/old cells (elongation and maturation zone) respective to the QC. The expression levels of the stem cell markers *TMO7* and *PLT2* are also shown. Data obtained from the Arabidopsis Plant eFP Viewer (bar.utoronto.ca).

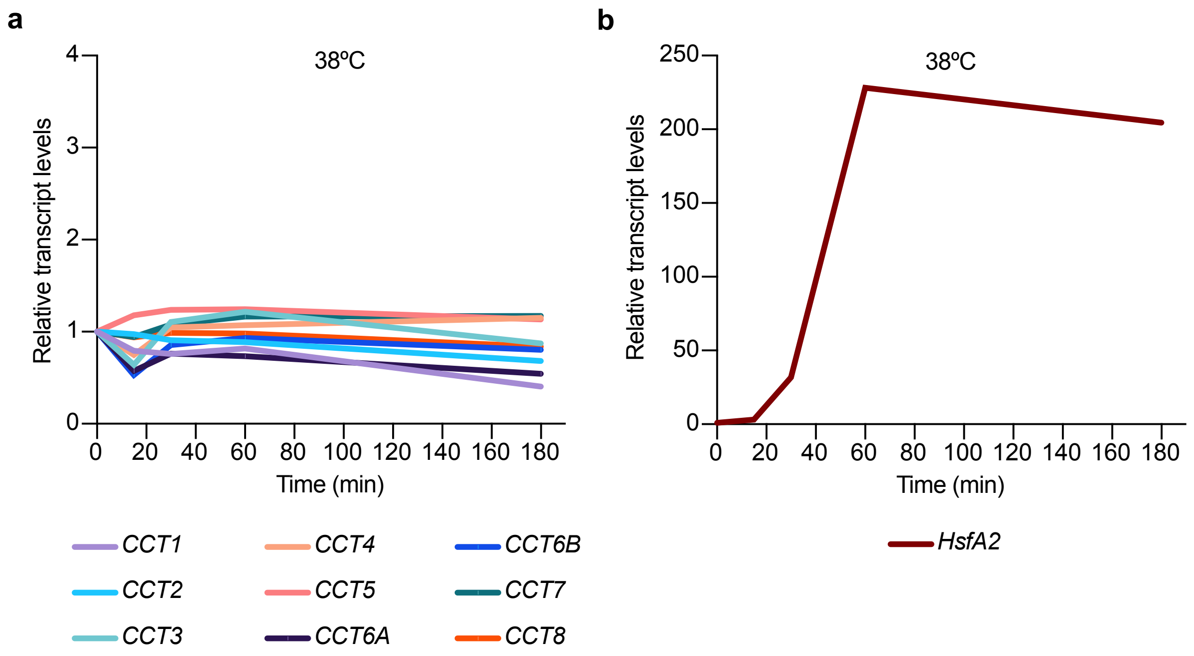


**Supplementary Figure 6. *CCT* subunits do not increase after heat shock treatment. a,** Relative transcript levels of genes encoding CCT subunits after heat shock. Arabidopsis plants grown under long-day conditions were transferred from control condition at 24°C to an incubator at 38°C. Samples from roots were recollected and analyzed at 0, 15, 30, 60, 180 min after transfer to heat stress. **b,** Expression levels of the heat shock regulator *HsfA2* increases rapidly after heat stress exposure. Data obtained from the Arabidopsis Plant eFP Viewer (bar.utoronto.ca).

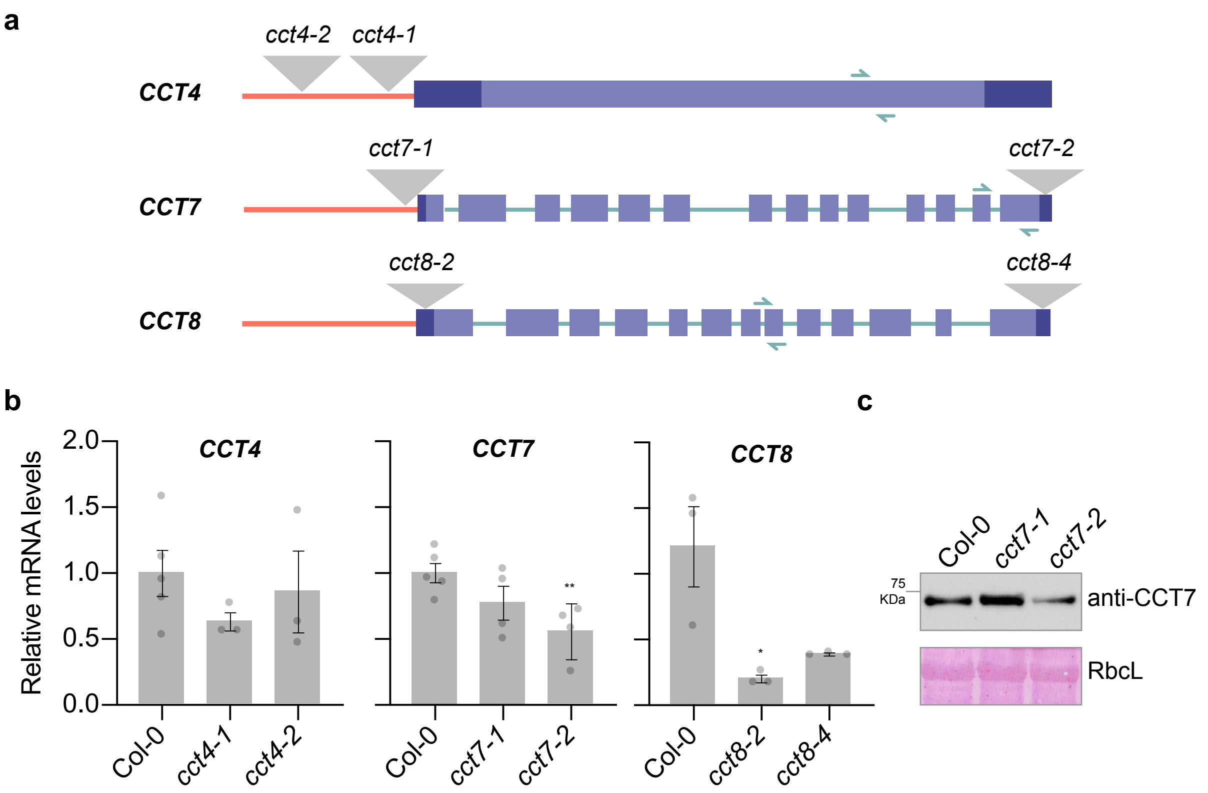


**Supplementary Figure 7. Characterization of *cct* mutant lines. a,** Schematic representation of the indicated *CCT* genes according to The Arabidopsis Information Resource (TAIR). The promoter region is represented as an orange line, the protein coding sequence (corresponding to the exons) is shown in purple, while the 3' and 5' untranslated regions are shown in dark purple. The approximate T-DNA insertions are represented with gray triangles. Green arrows indicate the position of the qPCR primers used in **Supplementary Fig. 7b**. The schemes are not-to-scale. **b,** qPCR analysis showing transcript levels of the indicated *CCT* genes in 6 DAG *cct* mutants compared to Col-0 wild-type (WT) (mean ± s.e.m of 3-4 independent experiments). Statistical comparisons were performed by two-tailed Student's *t*-test for unpaired samples. *P* value: **P*<0.05 ***P*<0.01. (C) Immunoblot with antibody against CCT7 of *cct7* mutants and Col-0 wild-type (15 DAG). Rubisco Large subunit (RbcL) is the loading control. Images are representative of 2 independent experiments.

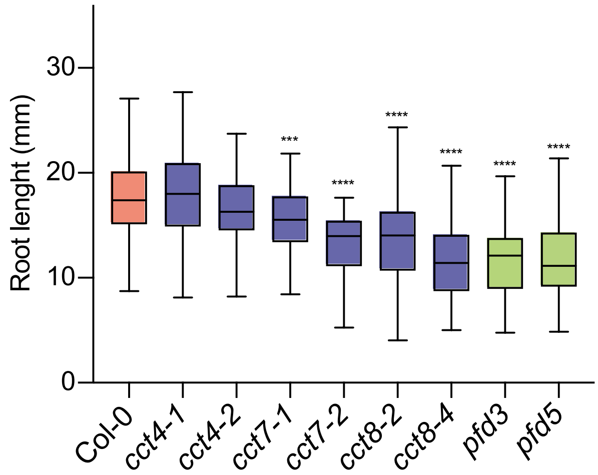


**Supplementary Figure 8. Root length screening assay of Arabidopsis chaperonin mutants.** Tukey box plots depicting root length in millimeters (mm) from 6 DAG plants. Horizontal line represents the median, box represents 25th and 75th percentiles. Col-0 (in pink), *cct* (in purple), and *pfd* (in green) mutants were analyzed (n= 25 seedlings from 3 independent experiments). Statistical analysis comparing Col-0 and mutants were performed with one-way ANOVA for multiple comparisons. *P* value: ***$P$<0.001, ****$P$<0.0001.

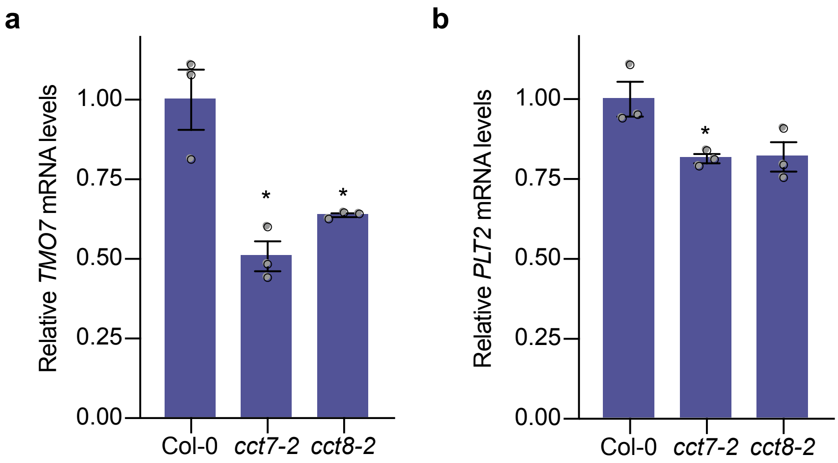


**Supplementary Figure 9. *cct* mutations result in decreased expression of meristem maintenance markers.** qPCR analysis showing transcript levels of *TMO7* (**a**) and *PLT2* (**b**) genes in 6 DAG seedlings relative to Col-0 (WT). Graphs represent the mean ± s.e.m of 3 independent experiments. Statistical comparisons were made by two-tailed Student's *t*-test for unpaired samples. *P* value: **P*<0.05, ***P*<0.01.

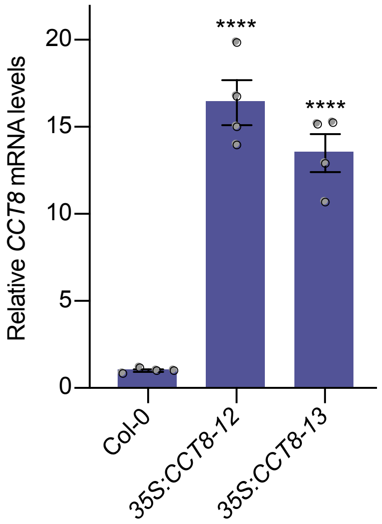


**Supplementary Figure 10. CCT8 mRNA levels in transgenic *35S:CCT8* hygromycin-resistant Arabidopsis lines.** qPCR analysis of *CCT8* transcript levels relative to age-matched WT seedlings (6 DAG). Graph represents the mean ± s.e.m of 4 independent experiments. The statistical comparisons were made by two-tailed Student's *t* -test for unpaired samples. *P* value: ****$P$<0.0001.

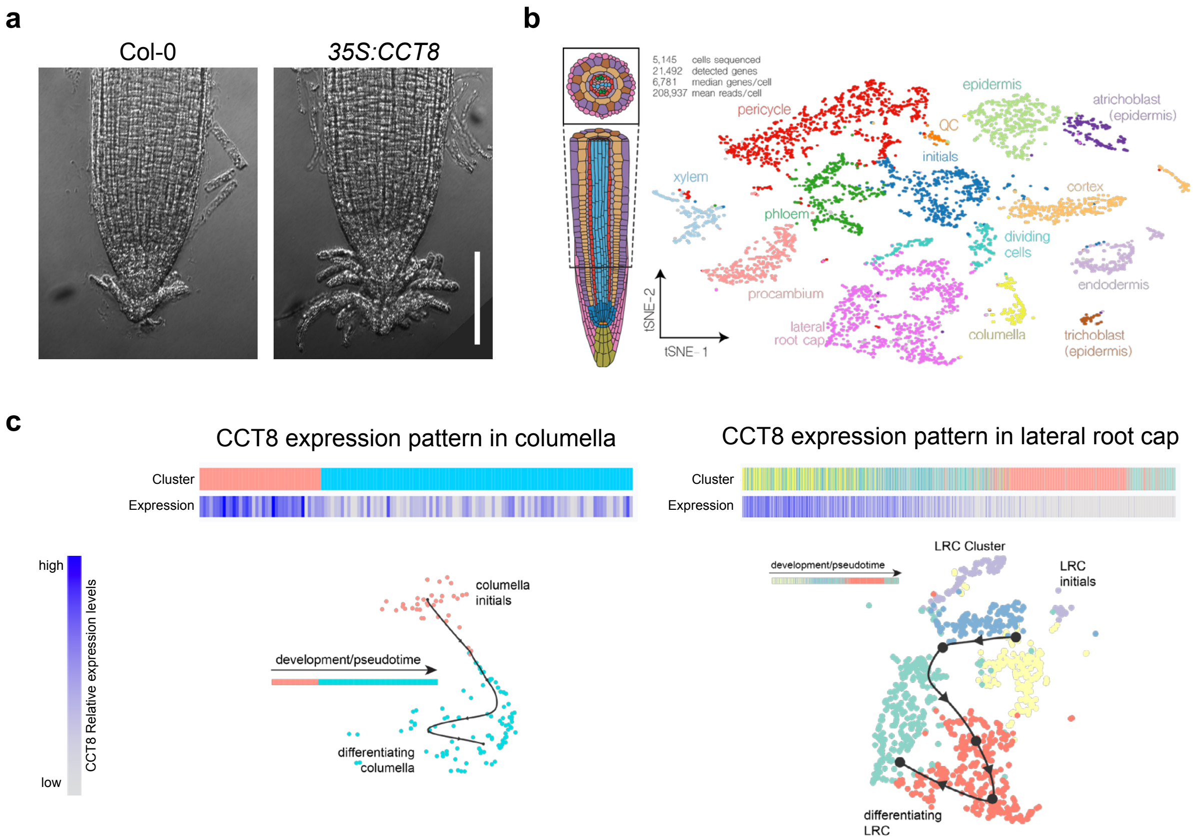


**Supplementary Figure 11. Cell-type expression pattern of *CCT8* in the columella and lateral root cap.** **a,** Representative images of the sloughing lateral root cap of Col-0 WT and *35S:CCT8* grown on plates supplemented with 15 µM MG-132. **b,** Color-coded tSNE plot showing the classification of 5145 high-quality (UMI count >17,290) cells into distinct cell identities corresponding to the schematic representation of the root meristem on the left. **c,** *CCT8* expression in the columella and lateral root cap decreases during development. Data were obtained from the on-line tool for visualization of cell type specific expression patterns of individual genes along developmental trajectories (http://bioit3.irc.ugent.be/plant-sc-atlas/).
